## Supplementary material for "Translocation of an arctic seashore plant reveals signs of maladaptation to altered climatic conditions"

Supplementary material for Hällfors et al. (submitted manuscript): **Translocation of an arctic seashore plant reveals signs of maladaptation to altered climatic conditions**

Text S1
Tables S1-S8
Figures S1-S6

**Text S1. Measuring local adaptation**

In their extensive guide to measuring local adaptation, Blanquart, Kaltz, Nuismer, and Gandon (2013) reviewed three conceptually differing definitions of local adaptation (LA) and the subsequent interpretations that can be drawn from experiments based on these. They also highlighted the fact that the estimation of local adaptation is sensitive to the definition used to characterise LA at the population level (Blanquart, Kaltz, Nuismer, & Gandon 2013). One of them (i) gives a general estimate on how local genotypes are adapted to their home environments and other two (ii and iii, reviewed in Kawecki and Ebert 2004) yield information on variation within the species concerned (i.e. genotype specific local adaptation). For an in-depth discussion on these concepts and their implications for measuring local adaptation, please see Kawecki and Ebert (2004) and Blanquart, Kaltz, Nuismer, & Gandon (2013).

i. The sympatric-allopatric contrast (henceforth called ΔSA) is the most straightforward way of estimating LA. In this approach the difference is calculated between the average fitness of genotypes in their home environment and genotypes in their non-home environment. In the case of this study ΔSA for the *subspecies as whole* is the difference between the mean fitness of both varieties in their home environments and that in the test (away) environment:

$$\Delta SA = \frac{p_{NH} + p_{SH}}{2} - \frac{p_{NA} + p_{SA}}{2}$$

ii. The home vs. away approach (henceforth called HA) can conceptually be understood as describing the *home environment quality for each tested genotype*, and is measured by the difference between the fitness of a genotype in its home environment and its fitness in all other tested environments (away). In our study, the Home vs. Away metric (HA) for each *variety* is the difference between the fitness of each variety in its home environment and that in its away environment.

$$HA_S = p_{SH} - p_{SA}$$

$$HA_N = p_{NH} - p_{NA}$$

iii. The local vs. foreign approach (henceforth called LF) can, in turn, be understood conceptually as describing the *genotype quality for each tested home environment*, and is measured by the difference between the fitness of a genotype in its home environment and the mean fitness of all other genotypes when exposed to the same environment. In our study the Local vs. Foreign metric (LF) for each *location* is the difference between the fitness of the local variety and that of the foreign variety.

$$LF_S = p_{SH} - p_{NA}$$

$$LF_N = p_{NH} - p_{SA}$$

**TABLE S1.** Seed collection information for populations used in experiments. The distance between sampled individuals in a seed sampling sites was approximately between 0.5 and 150 meters. The mean distance between the seed sampling sites of the two varieties was 533.5 kilometers. Herbarium specimens are placed in the herbarium of Finnish Museum of Natural History, Helsinki (H). For each experimental garden, three seed-sampling sites (among A-K) were randomly chosen for both varieties to provide individuals (however, contingent on the availability of individuals; see manuscript). From each seed-sampling site (A-K), five unique individuals were randomly chosen to provide eight seeds for producing experimental plants (again, contingent on seed availability).

| Variety | Seed sampling site | Location | Municipality | Country | Latitude | Longitude |
|---|---|---|---|---|---|---|
| Southern; var. jokelae | A | Pauhunlahti | Ii | Finland | 65.38027778 | 25.29333333 |
| Southern; var. jokelae | B | Partalahti | Ii | Finland | 65.37583333 | 25.43722222 |
| Southern; var. jokelae | C | Praava | Ii | Finland | 65.28444444 | 25.30694444 |
| Southern; var. jokelae | D | Halosenniemi | Haukipudas | Finland | 65.2575 | 25.33305556 |
| Southern; var. jokelae | E | Villenniemi | Haukipudas | Finland | 65.22638889 | 25.28611111 |
| Northern; var. finmarchica | F | Svartaksla | Sør-Varanger | Norway | 69.71666667 | 30.13 |
| Northern; var. finmarchica | G | Lille Ropelv | Sør-Varanger | Norway | 69.77444444 | 30.19611111 |
| Northern; var. finmarchica | H | Jakobsnes | Sør-Varanger | Norway | 69.7275 | 30.12861111 |
| Northern; var. finmarchica | I | Lanabukt | Sør-Varanger | Norway | 69.74194444 | 30.46527778 |
| Northern; var. finmarchica | J | Storbukt | Sør-Varanger | Norway | 69.6825 | 30.45305556 |
| Northern; var. finmarchica | K | Jarfjordbotn | Sør-Varanger | Norway | 69.66361111 | 30.30055556 |

| Variety | Seed sampling site | Distance from the centre point of seed sampling sites of said variety (km) | Seed sampling date | N:o of individuals from which seeds were sampled | Herbarium specimen number | Individuals of the population were planted in the following gardens, based on random draw and availablity of individuals |
|---|---|---|---|---|---|---|
| Southern; var. jokelae | A | 8.57 | 27 Aug 2012 | 7 | 0140-0146 | Oulu |
| Southern; var. jokelae | B | 9.29 | 27 Aug 2012 | 57 | 0147-0202 | All gardens |
| Southern; var. jokelae | C | 2.54 | 27 Aug 2012 | 51 | 0203-0252 | All gardens |
| Southern; var. jokelae | D | 5.27 | 27 Aug 2012 | 10 | 0253-0262 | None of the gardens |
| Southern; var. jokelae | E | 8.98 | 27 Aug 2012 | 50 | 0263-0312 | All gardens |
| Northern; var. finmarchica | F | 5.74 | 1 Sept 2012 | 50 | 0313-0362 | Tartu, Oulu, and Svanvik |
| Northern; var. finmarchica | G | 7.06 | 1 Sept 2012 | 50 | 0363-0412 | Helsinki, Oulu, and Svanvik |
| Northern; var. finmarchica | H | 5.9 | 1 Sept 2012 | 50 | 0413-0462 | Tartu, Rauma, and Svanvik |
| Northern; var. finmarchica | I | 7.66 | 2 Sept 2012 | 50 | 0463-0512 | Helsinki, Rauma, and Oulu |
| Northern; var. finmarchica | J | 7.78 | 2 Sept 2012 | 25 | 0513-0537 | Rauma |
| Northern; var. finmarchica | K | 6.08 | 2 Sept 2012 | 50 | 0538-0587 | Tartu and Helsinki |

**TABLE S2.** Description of experimental gardens and dates of procedures on plant individuals planted in each experimental garden. Distance from Southern vs Northern variety refers to the distance in kilometers from the centre of all seed sampling sites of that variety (seed sampling sites presented in Table S1). Experiment supplementation refers to new plants added to the experiment during the summer of 2013 when the experiment was started (see manuscript).

| Experimental garden | Coordinates | Distance from center point of seed sampling sites of the Southern variety (km) | Distance from center point of seed sampling sites of the Northern variety (km) | Date of seed sowing | Date of transfer to main glass house | Date of fertilizing | Date of exp. setup | Date of exp. supplementation |
|---|---|---|---|---|---|---|---|---|
| Tartu | N58.38, E26.72 | 772.7 | 1271.3 | 18 March 2013 | 24 April 2013 | 29 April 2013 | 17 May 2013 | 22 August 2013 |
| Helsinki | N60.20, E24.95 | 567.7 | 1086.3 | 26-28 March 2013 | 2 May 2013 | 7 May 2013 | 21 May 2013 | 10 September 2013 |
| Rauma | N61.13, E21.50 | 502.1 | 1035.2 | 4-5 April 2013 | 10 May 2013 | 15 May 2013 | 29-30 May 2013 | 2 September 2013 |
| Oulu | N65.06, E25.47 | 27.5 | 556.5 | 25-26 April 2013 | 1 June 2013 | 6 June 2015 | 17-18 June 2013 | 5 September 2013 |
| Svanvik | N69.45, E30.04 | 503.1 | 30.7 | 2 May 2013 | 7 June 2013 | 12 June 2013 | 27-28 June 2013 | 3 September 2013 |

**TABLE S3.** Number and percent of surviving individuals per year and from previous year in each garden. Also see Fig 3 in the main manuscript for a graphical representation.

| Garden | Variety | Planted individuals in 2013 | Survival percentage from 2013-2014 | Surviving individuals in 2014 | Survival percentage from 2014-2015 | Surviving individuals in 2015 | Survival percentage from 2015-2016 | Surviving individuals in 2016 and decreas from 2015 | Percent surviving by 2016 per variety, compared to number of individuals planted in 2013 | Percent surviving by 2016 across all individuals, compared to number of individuals planted in 2013 |
|---|---|---|---|---|---|---|---|---|---|---|
| Tartu | Southern | 80 | 25.0 | 20 | 60.0 | 12 | 66.7 | 8 | 10.0 | 5.8 |
| | Northern | 75 | 10.7 | 8 | 75.0 | 6 | 16.7 | 1 | 1.3 | |
| Helsinki | Southern | 42 | 61.9 | 26 | 65.4 | 17 | 76.5 | 13 | 31.0 | 15.9 |
| | Northern | 65 | 16.9 | 11 | 45.5 | 5 | 80.0 | 4 | 6.2 | |
| Rauma | Southern | 61 | 39.3 | 24 | 70.8 | 17 | 82.4 | 14 | 23.0 | 11.8 |
| | Northern | 66 | 10.6 | 7 | 28.6 | 2 | 50.0 | 1 | 1.5 | |
| Oulu | Southern | 49 | 55.1 | 27 | 77.8 | 21 | 57.1 | 12 | 24.5 | 17.2 |
| | Northern | 50 | 28.0 | 14 | 71.4 | 10 | 50.0 | 5 | 10.0 | |
| Svanvik | Southern | 51 | 78.4 | 40 | 95.0 | 38 | 94.7 | 36 | 70.6 | 51.0 |
| | Northern | 51 | 49.0 | 25 | 76.0 | 19 | 84.2 | 16 | 31.4 | |

**TABLE S4.** Estimates, standard errors, t-values or z-values, and p-values for the *Garden* aster model with only fixed effects (left) and including random effects (right). The intercept represents the survival in 2014 of the northern variety in Svanvik.

| Garden-model without random effects | | | | | Garden-model with random effects | | | |
|---|---|---|---|---|---|---|---|---|
| p-value | z value | Std. Error | Estimate | Variable | Estimate | Std. Error | z value | p-value |
| <0.001 | -4.024 | 0.155 | -0.624 | (Intercept) | -0.550 | 0.171 | -3.225 | <0.01 |
| <0.001 | 3.718 | 0.329 | 1.223 | Survival 2015 | 1.212 | 0.328 | 3.695 | <0.001 |
| <0.001 | 6.547 | 0.263 | 1.720 | Survival 2016 | 1.638 | 0.263 | 6.228 | <0.001 |
| <0.001 | -8.066 | 0.352 | -2.843 | Flowering 2014 | -2.862 | 0.349 | -8.190 | <0.001 |
| <0.001 | -11.841 | 0.359 | -4.253 | Flowering 2015 | -4.241 | 0.356 | -11.904 | <0.001 |
| <0.001 | -8.048 | 0.373 | -3.001 | Flowering 2016 | -3.029 | 0.371 | -8.175 | <0.001 |
| <0.001 | 10.430 | 0.170 | 1.778 | Flowering abundance 2014 | 1.650 | 0.172 | 9.571 | <0.001 |
| <0.001 | 13.962 | 0.163 | 2.273 | Flowering abundance 2015 | 2.142 | 0.165 | 12.966 | <0.001 |
| <0.001 | 12.190 | 0.167 | 2.040 | Flowering abundance 2016 | 1.907 | 0.170 | 11.229 | <0.001 |
| <0.001 | 6.333 | 0.015 | 0.092 | Original size | 0.132 | 0.020 | 6.708 | <0.001 |
| <0.001 | 3.492 | 0.034 | 0.119 | Southern variety | 0.130 | 0.045 | 2.870 | <0.01 |
| <0.05 | -2.546 | 0.079 | -0.200 | Oulu | -0.199 | 0.125 | -1.596 | 0.100 |
| <0.01 | -3.133 | 0.142 | -0.445 | Rauma | -0.498 | 0.175 | -2.850 | <0.01 |
| <0.01 | -3.132 | 0.100 | -0.312 | Helsinki | -0.344 | 0.140 | -2.455 | <0.05 |
| <0.01 | -3.303 | 0.104 | -0.343 | Tartu | -0.375 | 0.144 | -2.606 | <0.01 |
| 0.500 | 0.689 | 0.086 | 0.060 | Southern:Oulu | 0.070 | 0.093 | 0.755 | 0.500 |
| 0.080 | 1.737 | 0.146 | 0.254 | Southern:Rauma | 0.247 | 0.151 | 1.637 | 0.100 |
| <0.01 | 2.591 | 0.104 | 0.270 | Southern:Helsinki | 0.279 | 0.109 | 2.558 | <0.05 |
| 0.200 | 1.336 | 0.109 | 0.146 | Southern:Tartu | 0.098 | 0.113 | 0.868 | 0.340 |
| -- | -- | -- | -- | Plot | 0.114 | 0.027 | 4.142 | <0.001 |
| -- | -- | -- | -- | Seed sampling site | 0.033 | 0.024 | 1.395 | 0.080 |

**TABLE S5.** Estimates, standard errors, t-values or z-values, and p-values for the *Annual temperature* aster model with only fixed effects (left) and including random effects (right). The intercept represents the survival in 2014 of the northern variety in Svanvik.

| Temperature-model without random effects | | | | | Temperature-model with random effects | | | |
|---|---|---|---|---|---|---|---|---|
| p-value | z value | Std. Error | Estimate | Variable | Estimate | Std. Error | z value | p-value |
| <0.05 | -3.021 | 0.165 | -0.499 | (Intercept) | -0.397 | 0.195 | -2.031 | 0.042 |
| <0.001 | 3.693 | 0.329 | 1.216 | Survival 2015 | 1.207 | 0.328 | 3.681 | <0.001 |
| <0.001 | 6.622 | 0.263 | 1.739 | Survival 2016 | 1.636 | 0.263 | 6.219 | <0.001 |
| <0.001 | -8.045 | 0.353 | -2.843 | Flowering 2014 | -2.868 | 0.350 | -8.205 | <0.001 |
| <0.001 | -11.865 | 0.360 | -4.274 | Flowering 2015 | -4.251 | 0.356 | -11.932 | <0.001 |
| <0.001 | -8.039 | 0.374 | -3.005 | Flowering 2016 | -3.036 | 0.371 | -8.192 | <0.001 |
| <0.001 | 10.643 | 0.170 | 1.813 | Flowering abundance 2014 | 1.651 | 0.172 | 9.572 | <0.001 |
| <0.001 | 14.231 | 0.163 | 2.312 | Flowering abundance 2015 | 2.143 | 0.165 | 12.970 | <0.001 |
| <0.001 | 12.451 | 0.167 | 2.079 | Flowering abundance 2016 | 1.908 | 0.170 | 11.233 | <0.001 |
| <0.001 | 5.923 | 0.012 | 0.072 | Original size | 0.126 | 0.019 | 6.618 | <0.001 |
| 0.980 | 0.029 | 0.055 | 0.002 | Southern variety | 0.038 | 0.066 | 0.580 | 0.562 |
| <0.001 | -5.199 | 0.014 | -0.075 | Temperature | -0.082 | 0.023 | -3.586 | <0.001 |
| <0.01 | 3.199 | 0.015 | 0.049 | Southern:Temperature | 0.044 | 0.016 | 2.806 | <0.01 |
| -- | -- | -- | -- | Plot | 0.132 | 0.031 | 4.290 | <0.001 |
| -- | -- | -- | -- | Seed sampling site | 0.034 | 0.023 | 1.521 | 0.060 |

**TABLE S6.** Predictions of plant performance for each variety in each experimental garden based on the Garden fixed-effects aster model. The same data is shown as a barplot in the main manuscript, Fig. 5(a).

| Garden | Variety | Predicted value | Std. Error |
|---|---|---|---|
| Svanvik | Northern | 1.467 | 0.387 |
| | Southern | 5.691 | 0.805 |
| Oulu | Northern | 0.243 | 0.147 |
| | Southern | 1.687 | 0.500 |
| Rauma | Northern | 0.039 | 0.036 |
| | Southern | 1.048 | 0.309 |
| Helsinki | Northern | 0.100 | 0.071 |
| | Southern | 4.101 | 0.879 |
| Tartu | Northern | 0.080 | 0.058 |
| | Southern | 0.986 | 0.259 |

**TABLE S7.** Predicted values and standard errors of plant performance for each variety per mean annual temperature based on the Temperature fixed-effects aster model. The same data is shown in a plot in the main manuscript, Fig. 5(b).

| Annual temp. | Variety | Predicted value | Std. Error |
|---|---|---|---|
| 2.00 | Northern | 1.530 | 0.403 |
| | Southern | 4.846 | 0.711 |
| 2.61 | Northern | 0.988 | 0.229 |
| | Southern | 4.254 | 0.567 |
| 3.22 | Northern | 0.644 | 0.144 |
| | Southern | 3.713 | 0.445 |
| 3.83 | Northern | 0.425 | 0.101 |
| | Southern | 3.224 | 0.352 |
| 4.44 | Northern | 0.286 | 0.077 |
| | Southern | 2.789 | 0.288 |
| 5.06 | Northern | 0.195 | 0.060 |
| | Southern | 2.405 | 0.252 |
| 5.67 | Northern | 0.136 | 0.047 |
| | Southern | 2.069 | 0.237 |
| 6.28 | Northern | 0.096 | 0.037 |
| | Southern | 1.777 | 0.232 |
| 6.89 | Northern | 0.068 | 0.029 |
| | Southern | 1.525 | 0.231 |
| 7.50 | Northern | 0.050 | 0.023 |
| | Southern | 1.309 | 0.229 |

**TABLE S8.** Local adaptation metrics calculated based on overall performance (Table S6) of the two varieties (LF= Local vs. Foreign; HA= Home vs. Away) and for both varieties (ΔSA= Sympatric-Allopatric contrast) in the reciprocal part of the experiment (Oulu and Svanvik).

| Variety | HA | LF | *ΔSA* |
|---|---|---|---|
| Southern | -0.71 | 0.26 | -0.25 |
| Northern | 0.22 | -0.75 | |

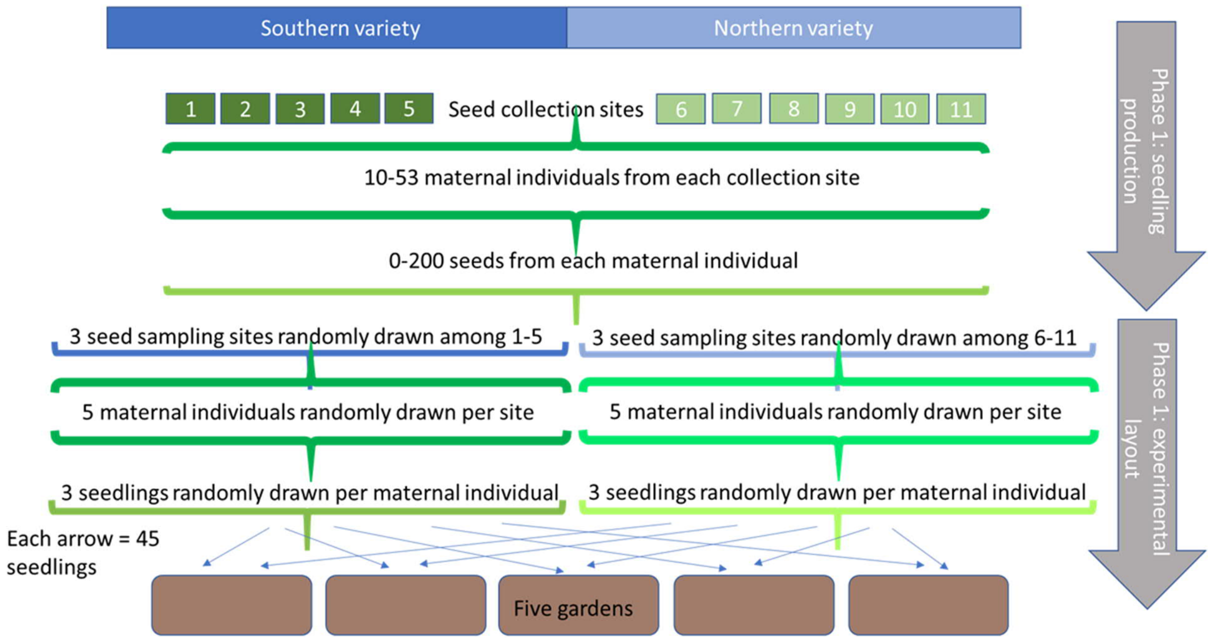


**FIGURE S1.** Framework for selecting seeds for growing experimental plants.

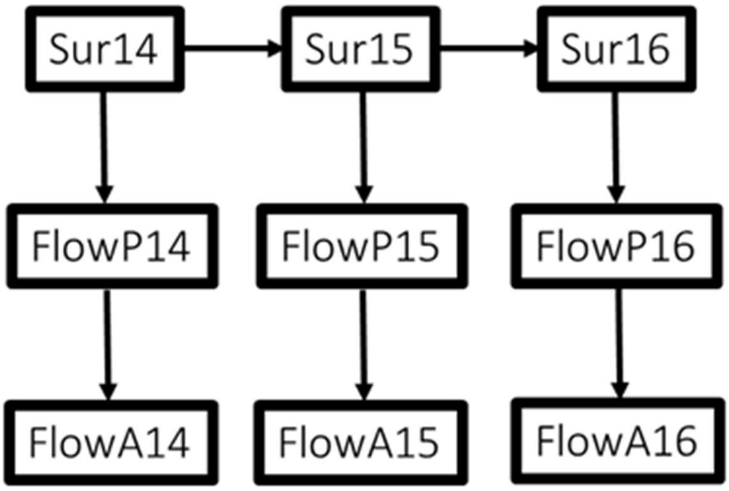


**FIGURE S2.** Graphical model for estimating overall performance using aster model. Performance measures are represented by the nodes and the arrows represent conditional distribution of the specified error. Specifically, the model included the probability of survival in each year (Sur14, Sur15, Sur16; 0 or 1; Bernoulli error distribution) that are conditional on survival in the previous year. Whether a plant flowered or not (FlowP14, FlowP15, FlowP16; 0 or 1; Bernoulli error distribution) is conditional on that it had survived in that year and in the previous year. The abundance of flowers per individual (FlowA14, FlowA15, FlowA16; count; zero-truncated poisson distribution of error) is conditional on that the individual flowered overall in that year and that it had survived in that year and in the previous year.

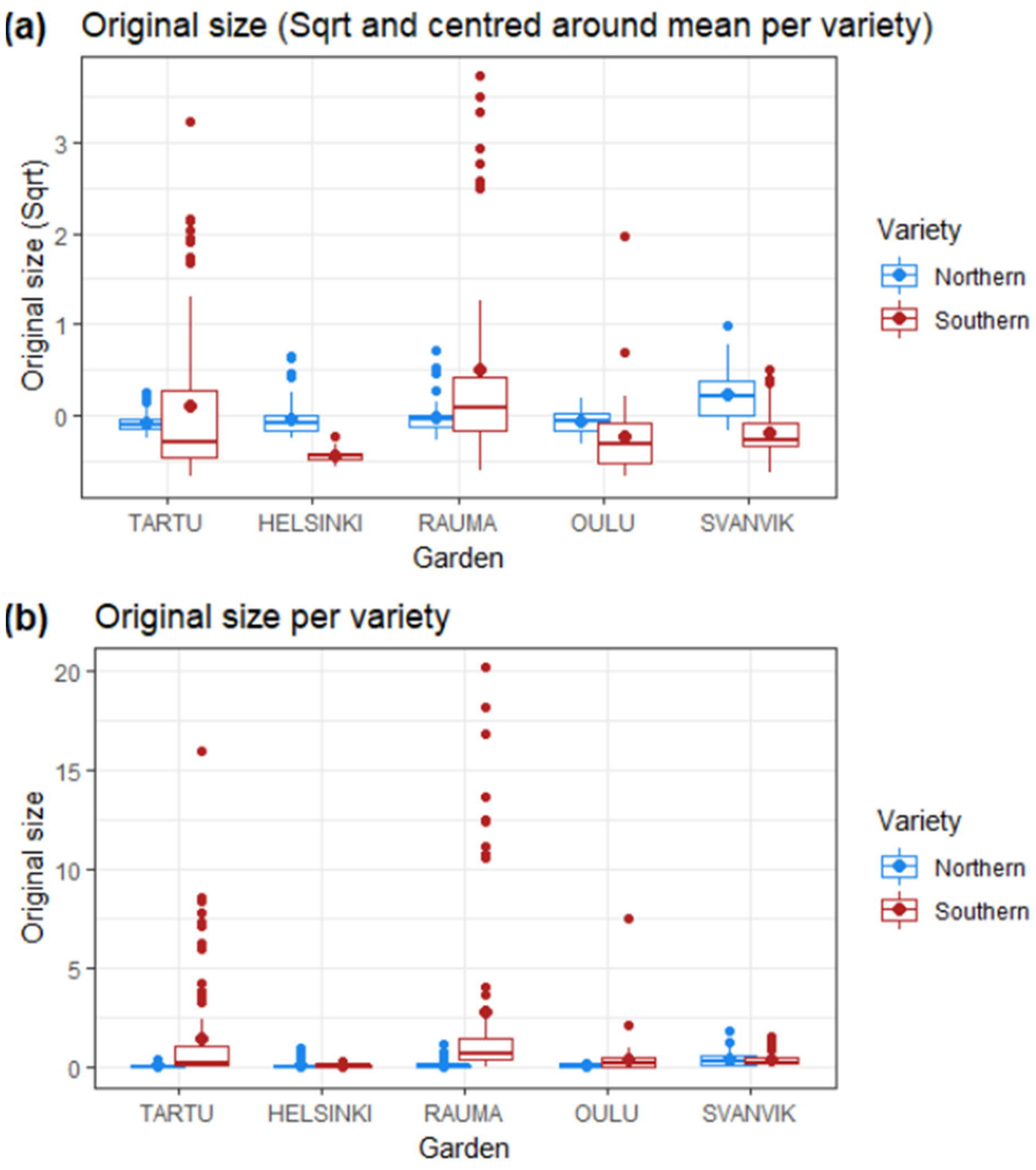


**FIGURE S3.** Original size of plants at planting in summer 2013. (a) Original size ($cm^2$) square root and centered around the mean of each variety. (b) Raw original size $cm^2$. For the analyzes, the square root transformed values were centered around the mean value for each variety separately (see Methods).

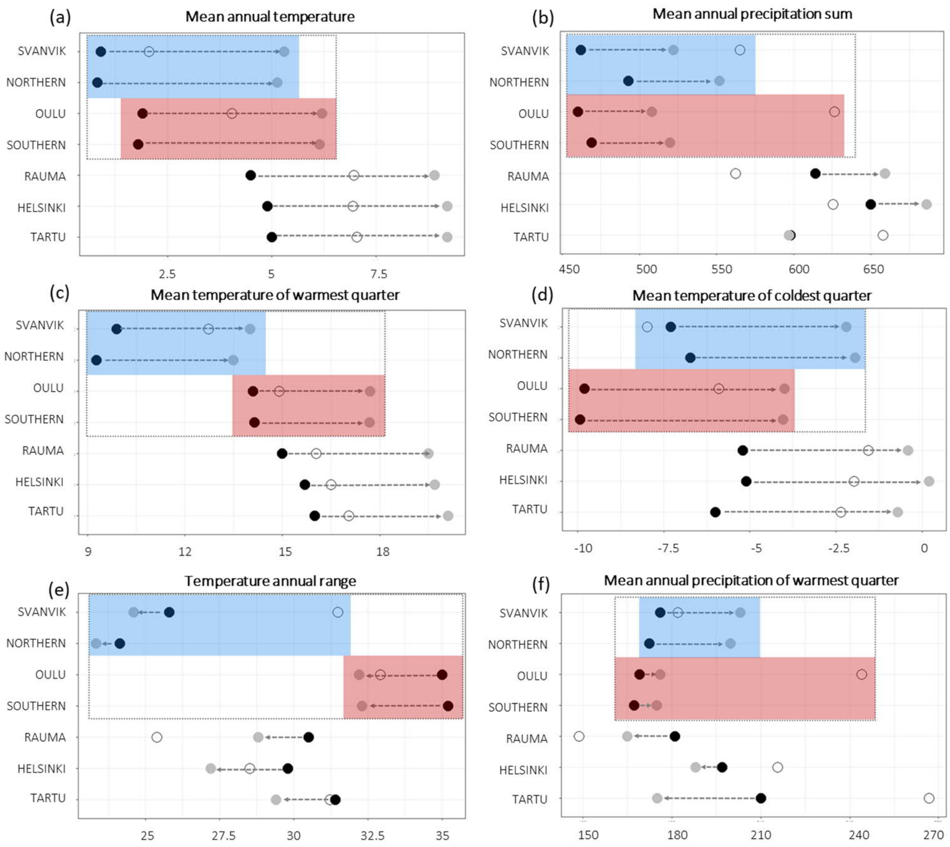


**FIGURE S4.** Climatic and weather conditions of seed sampling sites and experimental gardens. (a) Mean annual temperature (also shown in manuscript Figure 2); (b) mean annual precipitation sum; (c) mean temperature of the warmest quarter (mean temperature for the three months during which temperatures were highest in each year); (d) mean temperature of coldest quarter (mean temperature for the three months during which temperatures were lowest in that year); (e) temperature annual range (standard deviation * 100); and (e) mean annual precipitation of warmest quarter (the sum of precipitation for the three months during which temperatures were highest in that year). Black points show historic mean climatic conditions (1970-2000; 10 minutes resolution; Fick & Hijmans, 2017) and gray points show the future projections of mean climatic conditions (CMIP5 for 2050, 10 minutes resolution, HADGEM2-ES model; Fick & Hijmans, 2017) for each experimental garden and seed sampling site (Northern= seed sampling sites of the northern variety; Southern= seed sampling sites of the southern variety). For experimental gardens, open circles show mean climatic conditions during the experimental years 2013-2016. The experimental gardens and seed sampling sites within the species range is outlined by a dashed gray box, whereas the blue box indicates the sites within the range of the northern variety and the red box the sites within the range of the southern variety. All bioclim variables for the sites during historical, experimental, and future time periods are available at https://github.com/MariaHallfors/Primula-nutans-translocation.

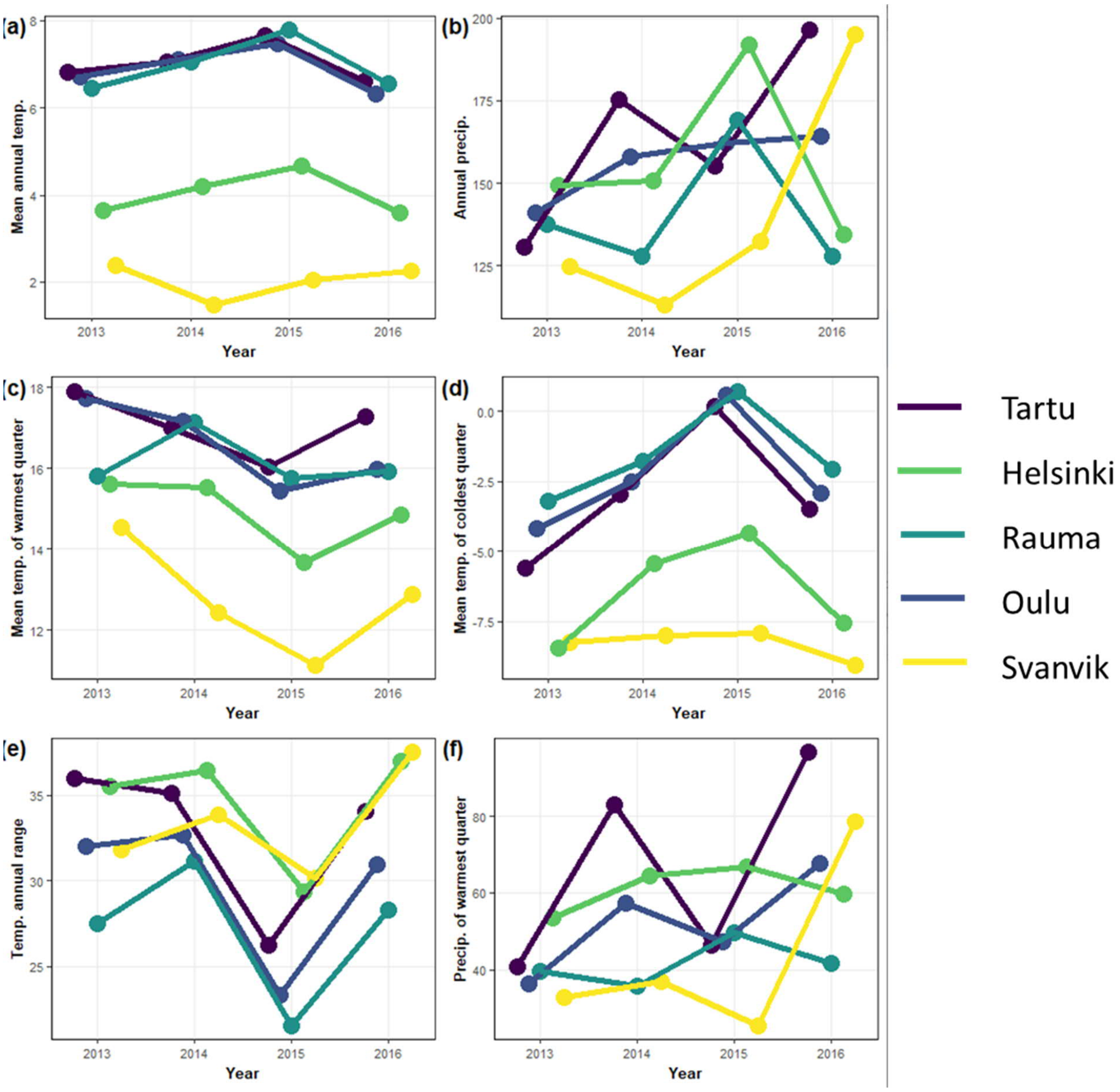


**FIGURE S5.** Annual weather conditions in the experimental gardens during the experimental years. (a) Mean annual temperature; (b) mean annual precipitation sum; (c) mean temperature of the warmest quarter (mean temperature for the three months during which temperatures were highest in each year); (d) mean temperature of coldest quarter (mean temperature for the three months during which temperatures were lowest in that year); (e) temperature annual range (standard deviation * 100); and (e) mean annual precipitation of warmest quarter (the sum of precipitation for the three months during which temperatures were highest in that year).

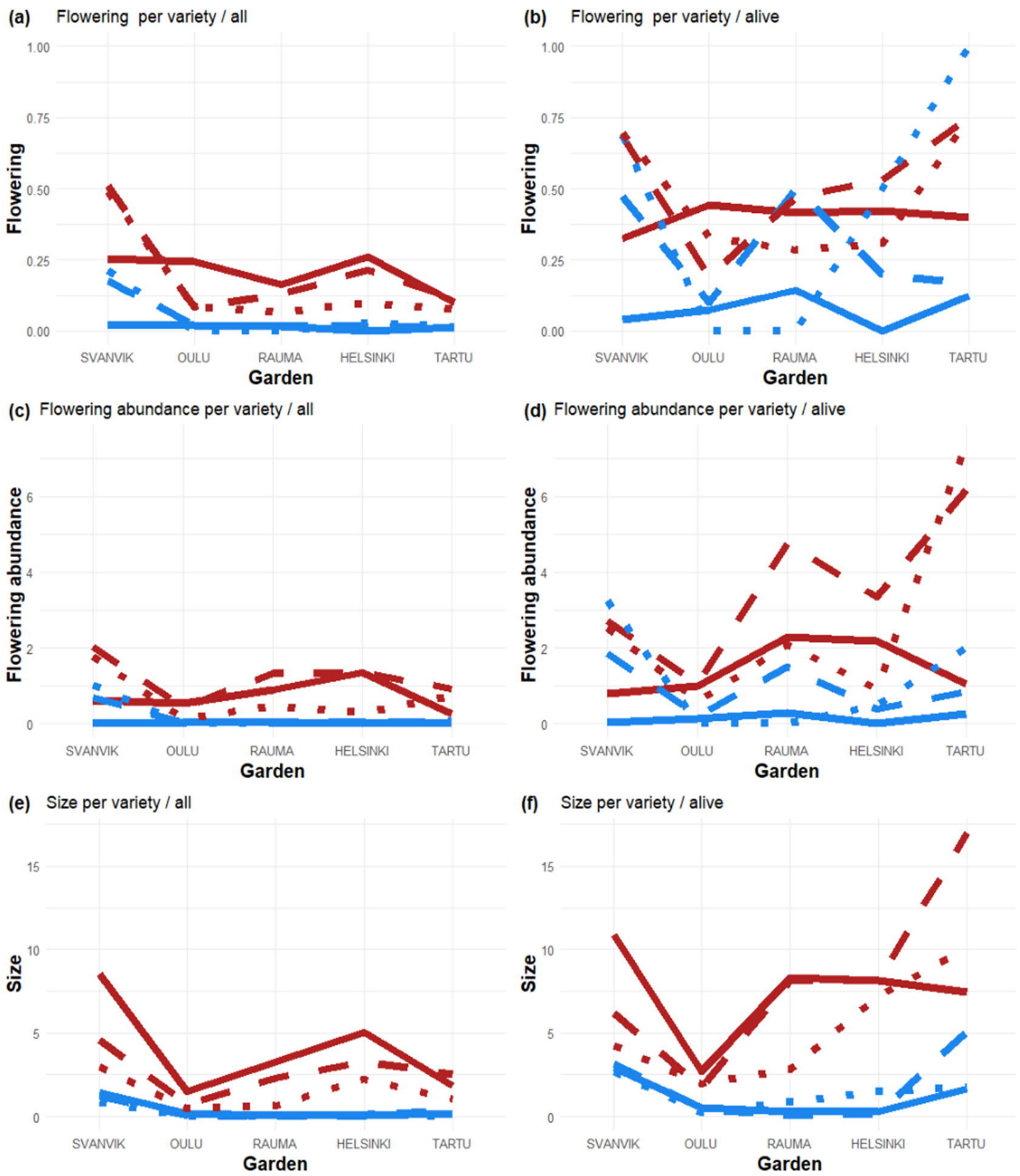


**FIGURE S6.** Raw data on performance measures per year, variety and experimental garden. Blue line= northern variety; red lines= southern variety; solid line= 2014; dashed line= 2015; dotted line= 2016. Panels in the left column shows yearly proportional means of performance measures (a) flower presence, (c) flowering abundance and (e) size ($cm^2$) averaged across all individuals were planted of each variety in each experimental garden, i.e. the performance measure is averaged across all planted individual (including dead ones which thus have a value of 0). Panels in the right column shows yearly absolute means of performance measures (b) flower presence, (d) flowering abundance and (f) size ($cm^2$) averaged across all individuals that were alive in the specific year of each variety in each experimental garden.
